# Caliber of zebrafish touch-sensory axons is dynamic in vivo

**DOI:** 10.1101/2024.12.04.626901

**Authors:** Kaitlin Ching, Alvaro Sagasti

## Abstract

Cell shape is crucial to cell function, particularly in neurons. The cross-sectional diameter, also known as caliber, of axons and dendrites is an important parameter of neuron shape, best appreciated for its influence on the speed of action potential propagation. Many studies of axon caliber focus on cell-wide regulation and assume that caliber is static. Here, we have characterized local variation and dynamics of axon caliber in vivo using the peripheral axons of zebrafish touch-sensing neurons at embryonic stages, prior to sex determination. To obtain absolute measurements of caliber in vivo, we paired sparse membrane labeling with super-resolution microscopy of neurons in live fish. We observed that axon segments had varicose or “pearled” morphologies, and thus vary in caliber along their length, consistent with reports from mammalian systems. Sister axon segments originating from the most proximal branch point in the axon arbor had average calibers that were uncorrelated with each other. Axon caliber also tapered across the branch point. Varicosities and caliber, overall, were dynamic on the timescale of minutes, and dynamicity changed over the course of development. By measuring the caliber of axons adjacent to dividing epithelial cells, we found that skin cell division is one aspect of the cellular microenvironment that may drive local differences and dynamics in axon caliber. Our findings support the possibility that spatial and temporal variation in axon caliber could significantly influence neuronal physiology.

**Significance Statement:** Axon caliber directly influences how quickly neurons send messages to other cells and likely plays a role in neurons’ overall health. In the peripheral nervous system, where neurons cover particularly long distances, cell shape can determine whether an animal successfully executes behaviors such as escape responses. We found that axon caliber can vary between locations within the same cell and is highly dynamic. Taking these variations into account may allow neuroscientists to better estimate transmission speeds for cells in neural circuits. We observed that axon caliber is distorted when nearby skin cells change shape. Thus, cells not classically considered part of the nervous system can also contribute to caliber dynamics, broadening our view of axon caliber determinants.

## Introduction

The shape of a neuron’s axons or dendrites is central to its function. A key characteristic of axon and dendrite shape is their cross-sectional diameter, also known as caliber. Axon caliber intrinsically influences the speed of action potential propagation (Kriz et al., 2000; Sakaguchi et al., 1993) and may also play a role in cell health and structural stability (Dollé et al., 2018; Perge et al., 2012).

Rohon-Beard (RB) neurons are a type of touch-sensing neuron that develops in early embryonic zebrafish and project skin-innervating peripheral arbors with branched architecture (Fig 1A) (Katz et al., 2021; Shorey et al., 2021; Wang et al., 2013). RB peripheral arbors are afferent axons that detect somatosensory stimuli in the skin and relay messages to interneurons via central axons in the spinal cord, thus promoting behavioral responses.

**Figure 1:**
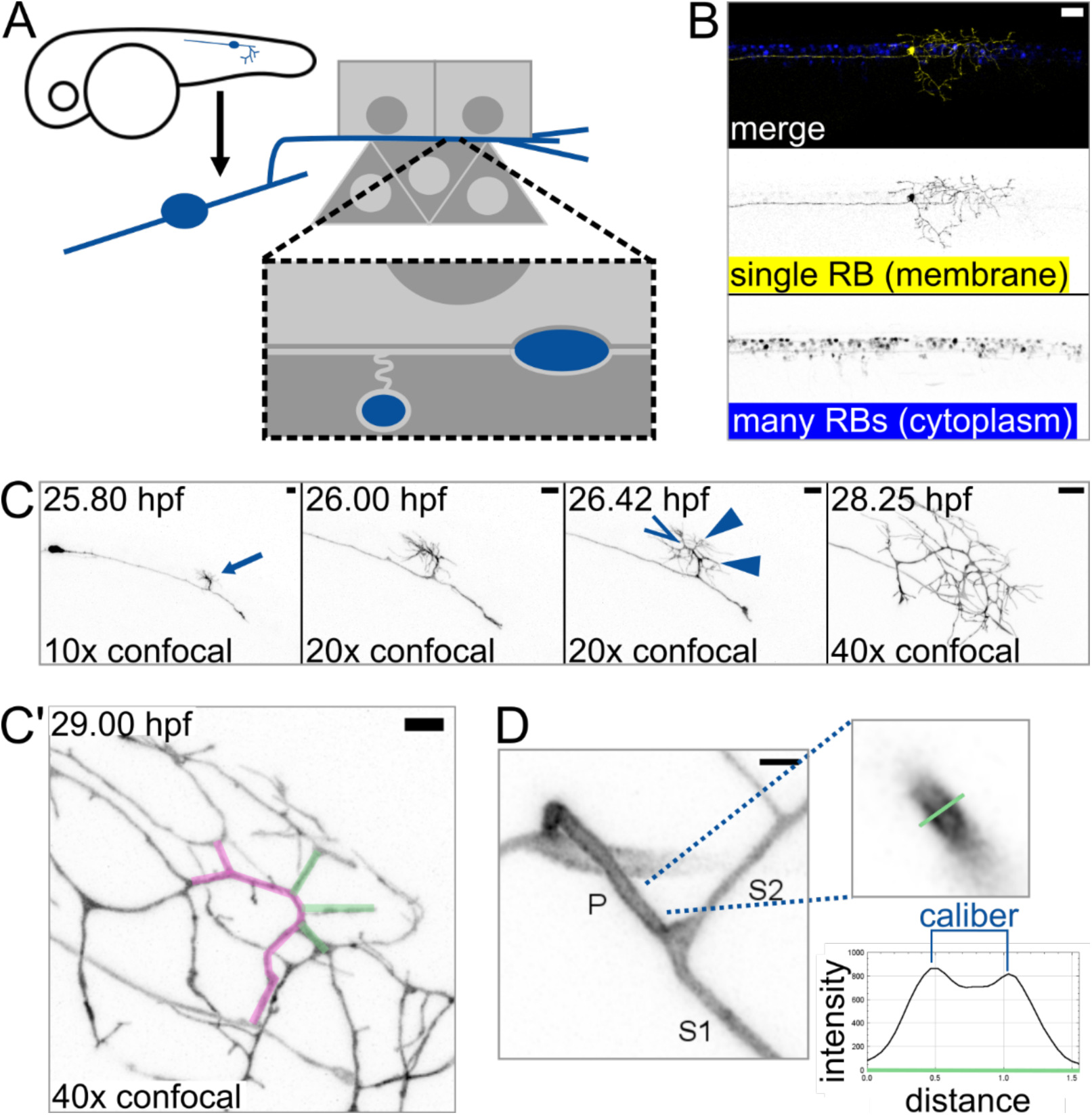
Axon branch development and caliber in sparsely-labeled Rohon-Beard (RB) neurons **(A)** Diagram depicting an RB neuron in an embryonic zebrafish. Light grey squares represent periderm (outermost) epithelial cells. Dark grey triangles represent basal epithelial cells, which ensheath some RB axons around 2 - 3 dpf. **(B)** Example of embryonic zebrafish tail with sparse labeling showing one RB neuron expressing EGFP-CAAX among many RB neurons expressing cytoplasmic DsRed. Scale bar: 50 µm. **(C)** Illustrative images taken from time lapse of RB neuron development. Proximal branch point formed by collateral sprouting in 10 out of 10 time lapses. Arrow: formation of main branch in peripheral axon arbor. 26.42 hpf panel shows multiple new branches that formed within minutes of each other. Open arrowhead: growth cone bifurcation event. Solid arrowheads: collateral sprouting events. Scale bars: 10 µm. **(C’)** Proximal region of the axon arbor shown in B. Magenta pseudo-colored axon shows the main branch, which branched by growth cone bifurcation. Green pseudo-colored axons show branches that formed by subsequent collateral sprouting. Scale bar: 5 µm. **(D)** Method used for measuring caliber. RB neurons expressing EGFP-CAAX were measured by line scan (green line). Caliber was scored as distance between intensity peaks, as illustrated in example plot. P: primary branch, which extends from spinal cord to skin, S1: thicker secondary branch, S2: thinner secondary branch. Scale bar: 2 µm.

**Figure 2:**
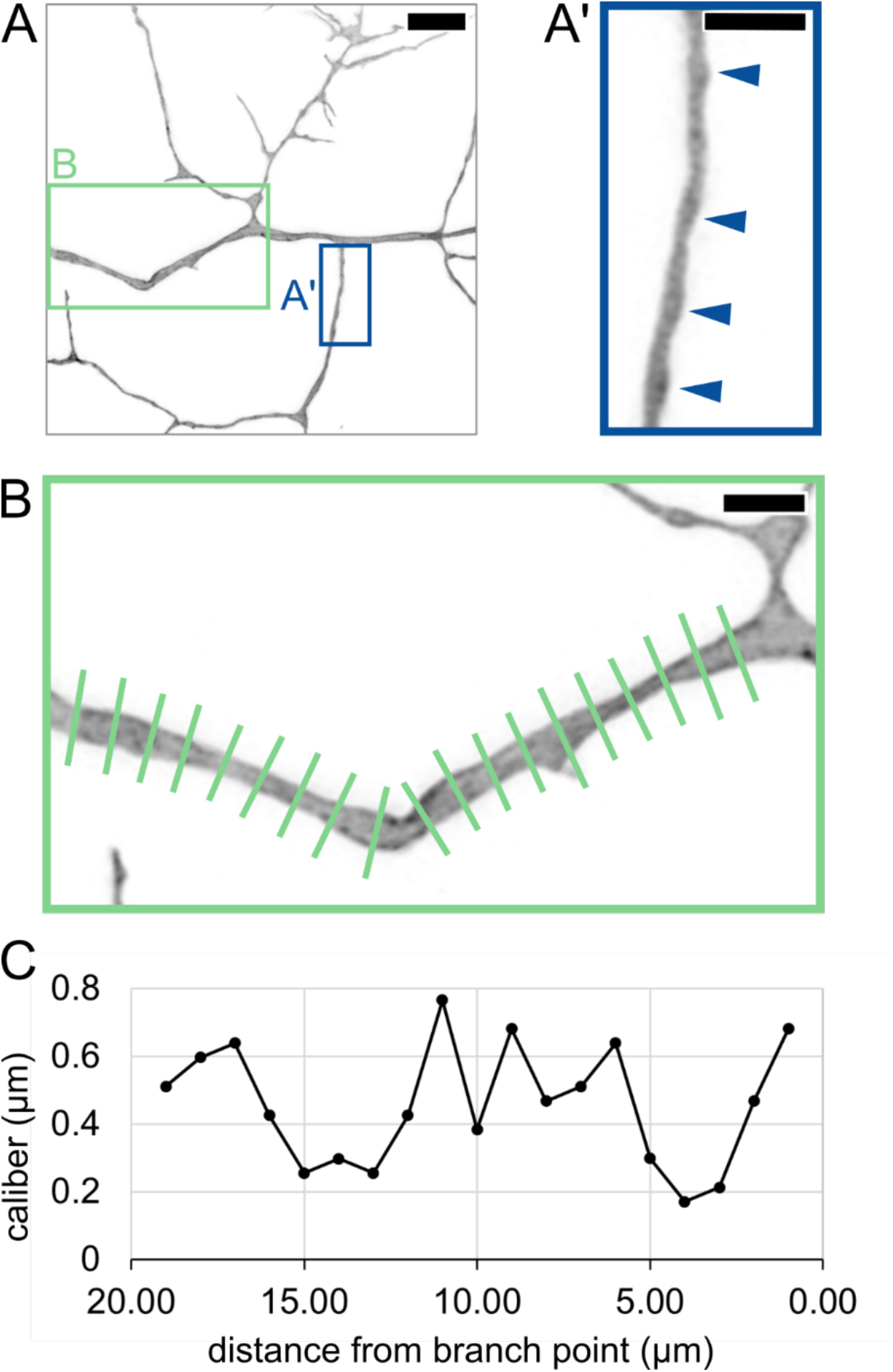
Lengthwise variation in axon caliber **(A)** Image of an RB axon arbor at 29.00 hpf. Blue box shows location of inset in A’. Green box shows location of inset in B. Scale bar: 5 µm. **(A’)** Inset showing axon branch with pearled morphology. Arrowheads: Thicker regions of axon, resembling “pearls” previously described in mammalian neurons. Scale bar: 2 µm. **(B)** Long axon segment measured by line scan at 1-µm increments. Scale bar: 2 µm. **(C)** Plot of caliber showing variation along the length of axon segment in panel B.

The significance of axon caliber in cell function has been described for about a century (Hursh, 1939), and it is widely regarded as a cell-wide, static trait despite reports of variations in axon or dendrite caliber over space and time. Axon caliber can vary along the length of axons due to inherent biophysical properties (Griswold et al., 2024), myelination (Seidl, 2014), and mechanical insult (Pan et al., 2024). Dendrite caliber is also known to taper across branch points (Craig and Banker, 1994). Axon caliber has been observed to change over development (Bomont, 2021; Seidl and Rubel, 2016), aging (Metzner et al., 2022), and in response to target size (Voyvodic, 1989a, 1989b) or activity (Nabel et al., 2024; Sinclair et al., 2017). These studies of caliber changes have primarily focused on changes to axon caliber over many days and assume that caliber undergoes little to no change on the timescale of hours. However, the cytoskeletal structures (Costa et al., 2020; Taylor et al., 2012; Wang et al., 2020) and cargoes (Costa et al., 2020; Greenberg et al., 1990; Wang et al., 2020) that can impact axon caliber are dynamic on the timescale of seconds to minutes, prompting us to ask how axon caliber and morphology change on shorter timescales.

Here, we characterized axon caliber in zebrafish RB neurons. We found that each single arbor, in vivo, can have a variety of axon calibers, varying within a segment and between sister segments. Axon calibers were dynamic on the timescale of minutes, and these dynamics changed over the developmental timescale of days. Finally, we found that the behavior of epithelial cells in the skin impacts sensory axon caliber, highlighting the complexities of the in vivo axon environment.

## Materials and Methods

### Fish care

Zebrafish (Danio rerio) adults were housed at 28.5°C with 13.5/10.5-hour light/dark cycles. Embryos were raised at 28.5°C in water containing 0.06 g/L Instant Ocean salt mix (Spectrum Brands, AA1-160P) and 0.05% methylene blue (ThermoFisher, 042771.AP).

### Injection for sparse labeling

To sparsely label RB neurons for observation of development and axon caliber, adult fish expressing Isl1[ss]:Gal4, UAS:dsRed (Sagasti et al., 2005) were crossed, and resulting embryos were allowed to develop for approximately one hour after fertilization. When embryos reached the 4-cell or 8-cell stage, a single cell in each embryo was injected with approximately 1-3 nL of plasmid containing the Gal4 effector, UAS:egfp-caax, diluted to 10-15 ng/µL in water, with or without phenol red (Sigma, P4758). After injection, embryos were raised as described above.

### Mounting embryos for microscopy

Unless otherwise indicated, embryos were immobilized on coverslips in 1.2% low-melt agarose dissolved in water (Fisher, BP165-25) and mounted with an equal volume of water containing 0.12 g/L Instant Ocean (Spectrum Brands, AA1-160P) and 0.4 mg/mL tricaine (MS-222, Western Chemical Inc.).

### Developmental time lapse imaging

Embryos injected for sparse labeling were sorted at approximately 20 hours post-fertilization (hpf) to find animals with a single cell expressing EGFP-CAAX in the tail. Selected embryos were immediately dechorionated using forceps and anesthetized in approximately 0.2 mg/mL tricaine (MS-222, Western Chemical Inc.). Embryos were mounted as described above. A cut was made in the agarose posterior to the tail using a fly pin probe to allow room for growth.

Embryos were imaged at 5-minute intervals on a ZEISS laser scanning microscope (LSM) 800 with a stage heater at 28°C. Magnification was increased as needed by changing objectives to see branching. The most proximal branch point at approximately 28 hpf was identified and visually tracked back to the point of formation within the time lapse for scoring.

### Line scan and symmetry analysis

Embryos were injected for sparse labeling, as described above. Embryos with a distinct EGFP-CAAX-positive neuron in the tail were dechorionated using forceps and anesthetized in approximately 0.2 mg/mL tricaine (MS-222, Western Chemical Inc.). For this experiment, embryos were immobilized in 1.1 – 1.2% low-melt agarose on No.1 coverslips (Epredia, 12460S) to improve available working distance, and water used in initial anesthetization was added before sealing with a glass slide. The primary axon segment was identified as the one that exited the spinal cord and entered the skin. The most proximal branch point was imaged on a ZEISS LSM 880 with Airyscan deconvolution at 27 – 31 hpf. If two neurons or two arbors were present in the same animal or cell, the more posterior one was scored. Line scan analysis was performed using the Line and Plot Profile tools in ImageJ (Schindelin et al., 2012) to find intensity values along a line perpendicular to the center axis of the axon. Axon caliber was measured as the distance between the two peaks on the plot that corresponded to the plasma membrane. For symmetry analysis, axons were measured 3, 4, and 5 µm away from the branch point and averaged to find the caliber of the given segment. The caliber of secondary branches was normalized by dividing by the caliber of the corresponding primary branch. We defined symmetry as S2/S1, which is equivalent to the normalized S2 caliber divided by the normalized S1 caliber.

### Caliber dynamics time lapse imaging

Embryos were injected for sparse labeling, as described above. Around 24 hpf, embryos were mounted as described above. The proximal region of the axon arbor was imaged on a ZEISS LSM 980, heated to approximately 28°C, and processed with Airyscan Joint Deconvolution at a maximum of five iterations. Time lapses with 5-minute intervals were started at 28 - 31 hpf and acquired until the axon drifted out of the field of view, typically one to two hours. After imaging, embryos were recovered from the mounting agarose using forceps and a probe and grown an additional 24 hours under normal conditions (described above), with the addition of 0.2 mM PTU (1-phenyl 2-thiourea, Sigma, P7629) in the water. At 52 - 55.5 hpf, embryos were mounted and imaged in the same manner as the first day. For scoring, axon segment calibers were measured 3 µm from a branch point at every frame of the time lapse, and segments used were each separated by at least one branch point. If the same segments could be identified on the second day, caliber was scored at the same locations for every frame of the time lapse.

### Dual color time lapse imaging of cell division

To observe axons near dividing cells, transgenic embryos expressing Isl1[ss]:lexA, lexAop:mRuby-caax and tp63:Gal4VP16; UAS:egfp-plcδ-ph (Rasmussen et al., 2015) were mounted as described above and imaged on a ZEISS LSM 980, heated to approximately 28°C, and processed with Airyscan Joint Deconvolution at a maximum of five iterations. Basal skin cells in a rounded state were identified by eye when embryos were 28 - 31 hpf and immediately imaged by time lapse with 5-minute intervals, until daughter basal cells appeared flattened for at least two time points. Axon caliber was measured by line scan analysis, as described above, at 1-µm intervals, excluding regions less than 3 µm from a branch point. Border to border length of axons was measured using the Line tool in ImageJ (Schindelin et al., 2012). Orthogonal projections were generated in ZEN Blue (ZEISS).

### Plotting

All dot plots were created in R. Code can be found at github.com/katieching/CaliberAnalysis. Histograms, dynamics, and pearling plots were created using Microsoft Excel.

### Experimental design and statistical analysis

For descriptive analyses, an appropriate size for the data set was chosen in advance, and images were collected in experimental batches until the predetermined size was exceeded. For statistical comparisons, a small data set was initially collected, and a power analysis was performed to determine the data set size needed to reach a p-value < 0.05. The data set was collected and analyzed before performing final statistical analyses. Data were excluded as necessary for technical reasons, namely image quality that was too poor for line scan analysis, segments where branch points were closer together than the distance indicated for the experiment (e.g. < 5 µm for symmetry analysis), images in which the primary branch caliber was below the limit of resolution for the system (Fig 3, 4), and axons in which > 20% of measurements at the 1 day post-fertilization (dpf) time point were below the limit of resolution for the system (Fig 5, 6). Significance of differences in paired data was evaluated by paired permutation test in R. Differences between measurements treated as populations were evaluated by Mann-Whitney test. Analyses performed in R can be found at github.com/katieching/CaliberAnalysis. The sex of animals included in the study was not considered because experiments occurred prior to sex determination.

**Figure 3:**
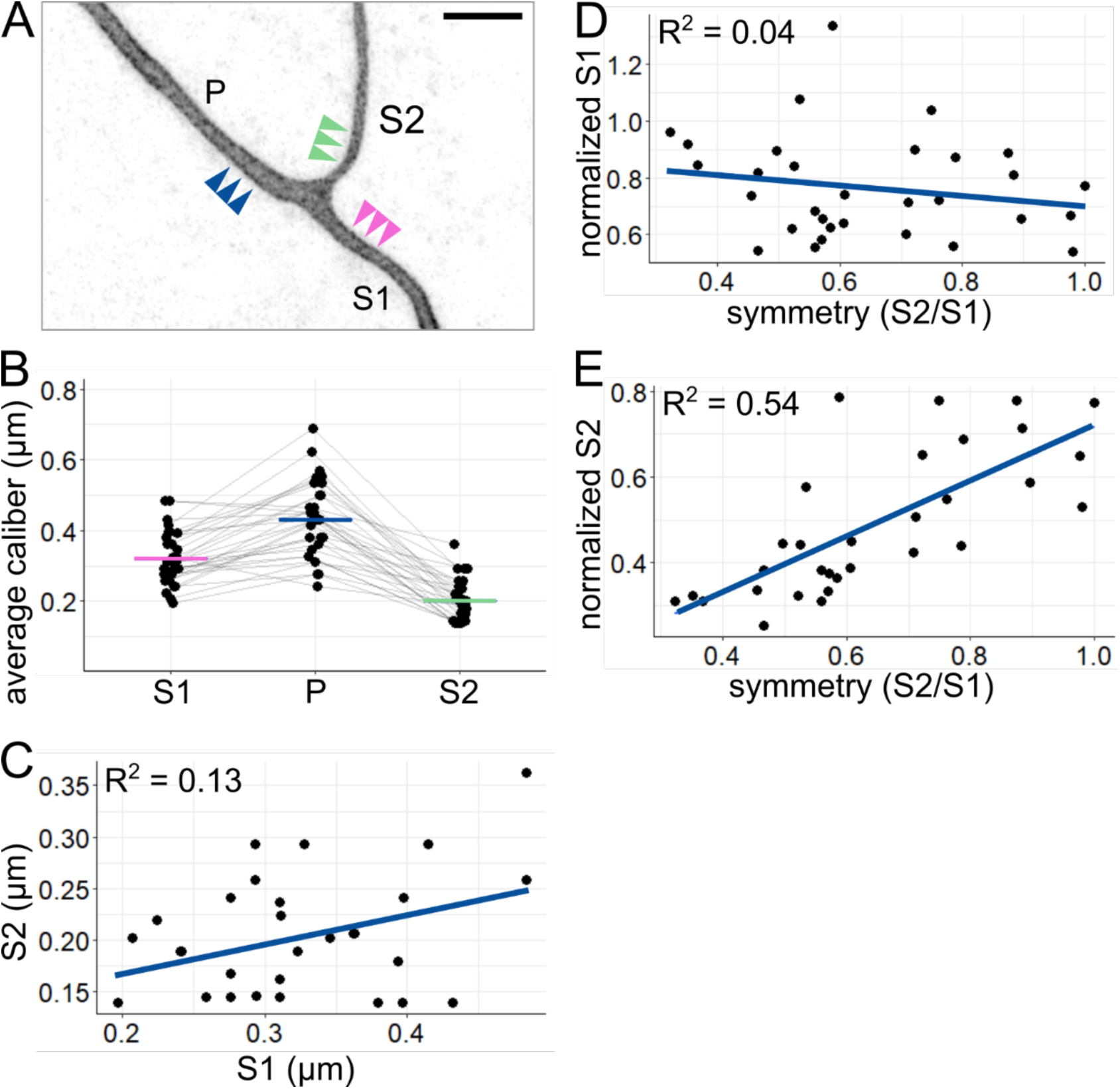
Comparison of axon segments within each arbor **(A)** Example image of proximal RB axon arbor. Arrowheads indicate approximate location of line scans for average caliber measurement. P: primary branch, S1: thicker secondary branch, S2: thinner secondary branch. Scale bar: 5 µm. **(B)** Plot of average axon calibers at the proximal branch point. Grey lines connect data points from the same neuron. Horizontal lines: mean values with colors corresponding to branch categories shown in panel A. n = 31 neurons. **(C)** Plot comparing caliber of sister axon segments shown in panel A. **(D)** Plot of branch symmetry versus normalized S1 caliber (S1/P). **(E)** Plot of branch symmetry versus normalized S2 caliber (S2/P). Blue lines in C, D, and E show linear regression.

**Figure 4:**
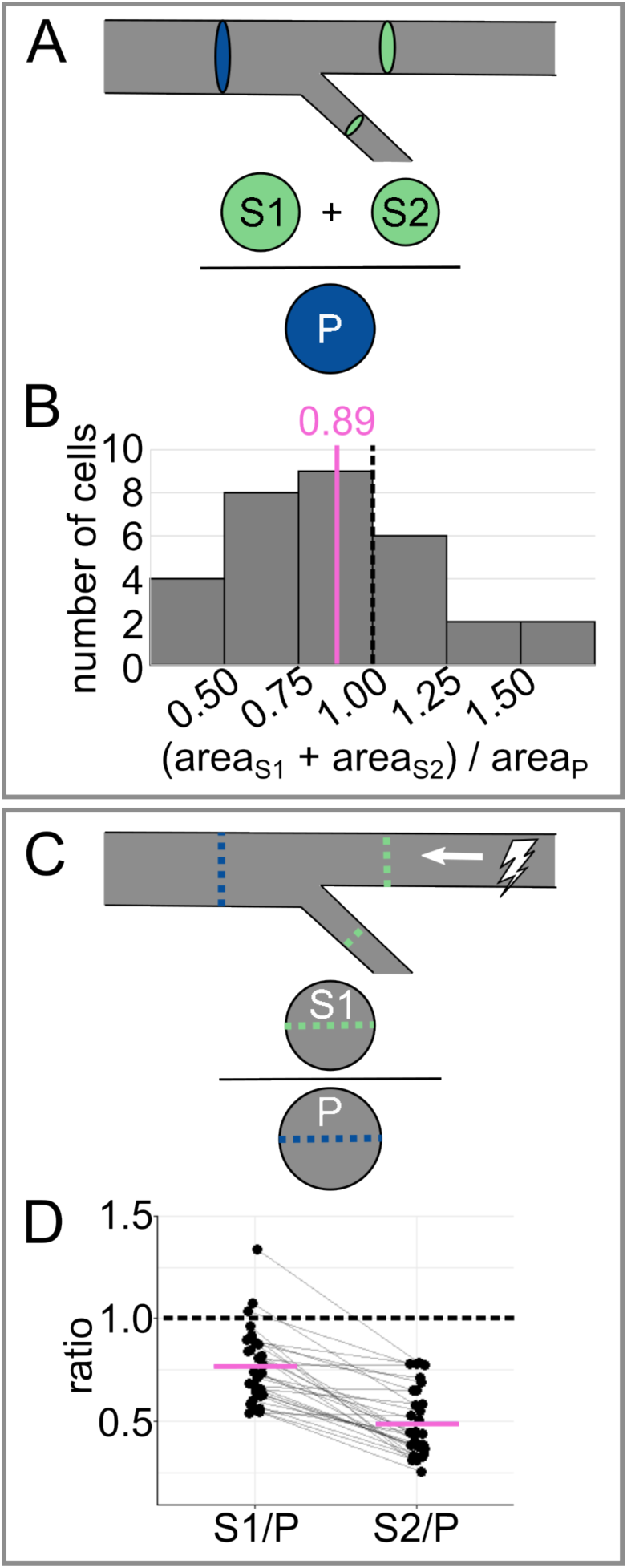
Analysis of tapering across the most proximal branch point **(A)** Diagram of tapering analysis in panel B. Combined cross-sectional area of secondaries (S1 and S2) is divided by the cross-sectional area of the primary (P) to evaluate tapering. **(B)** Histogram of cross-sectional area ratios for data shown in Figure 1. **(C)** Diagram of tapering analysis in panel D. Caliber of each secondary (S1 or S2) is divided by the caliber of the primary (P) to estimate the impact of tapering on action potential conduction velocity. **(D)** Plot of caliber ratios for data shown in Figure 1. Grey lines connect data points from the same neuron. In panels B and D, dotted black lines mark reference value 1.0 (no tapering), and magenta lines mark mean values.

**Figure 5:**
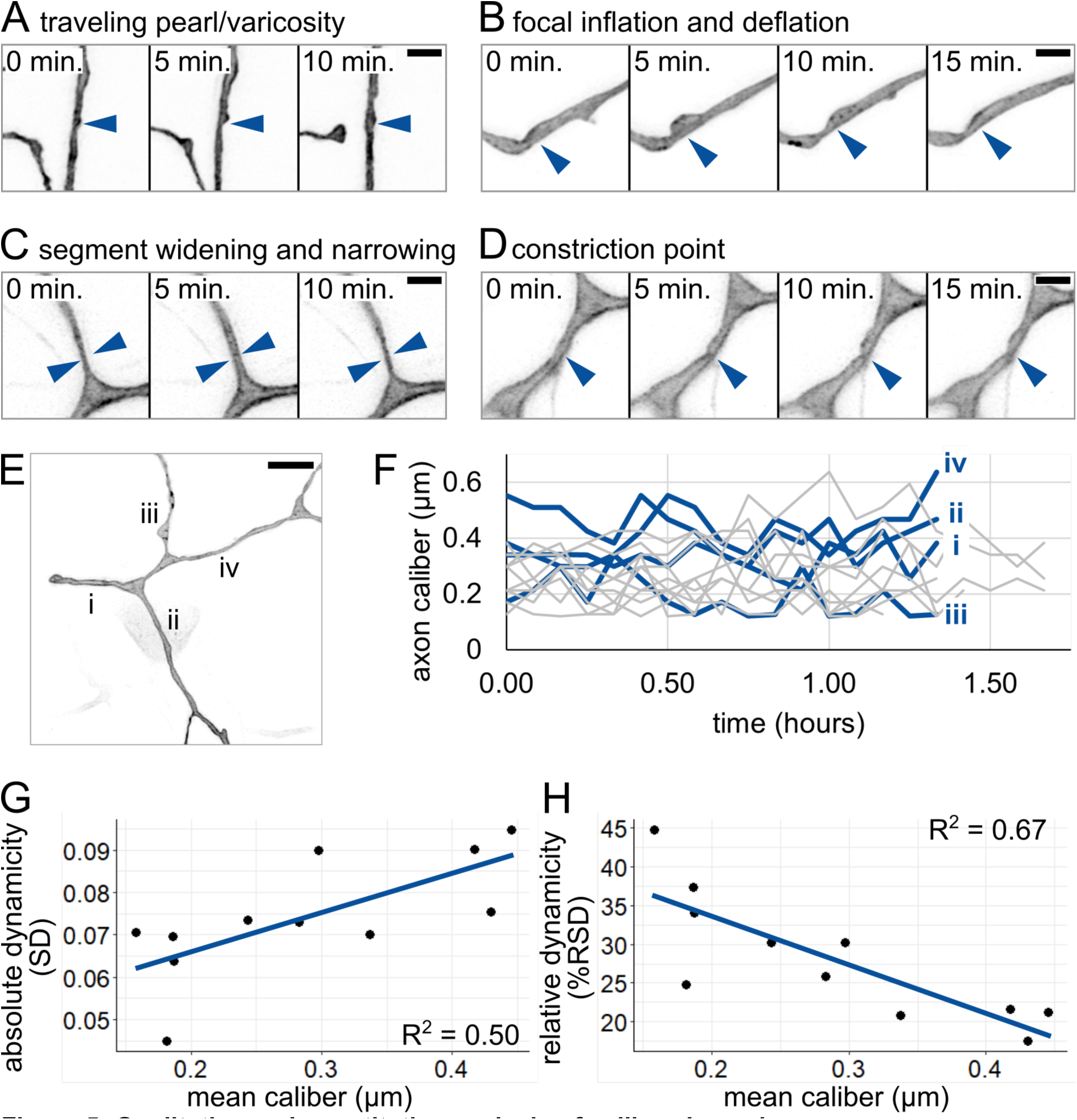
Qualitative and quantitative analysis of caliber dynamics **(A)** Example of possible traveling pearl. Short distance traveled allowed for acquisition despite 5-minute intervals. Arrowhead: initial location of pearl. **(B)** Example of focal inflation and deflation event. Puncta containing EGFP-CAAX accumulate during inflation and are seen exiting during deflation. Arrowhead: location of event. **(C)** Example of segment widening and narrowing. Arrowhead pairs denote width at t = 5 minutes. **(D)** Example of constriction point. Arrowhead: location of event. In panels A-D, scale bars: 2 µm. **(E)** Image of axon arbor in the proximal region. Different segments are labeled from proximal to distal: i-iv. Scale bar: 5 µm. **(F)** Plot of axon caliber dynamics, n = 11 axon segments, N = 5 fish. Blue lines: axon segments shown in panel E. Grey lines: all other axon segments measured. **(G)** Plot of mean caliber versus absolute dynamicity (SD) for all axons shown in panel F. Mean caliber is the average at the given location across all time points. **(H)** Plot of mean caliber versus relative dynamicity (%RSD = (SD / mean caliber) * 100) for all axons shown in panel F. In panels G and H, blue lines: linear regression.

**Figure 6:**
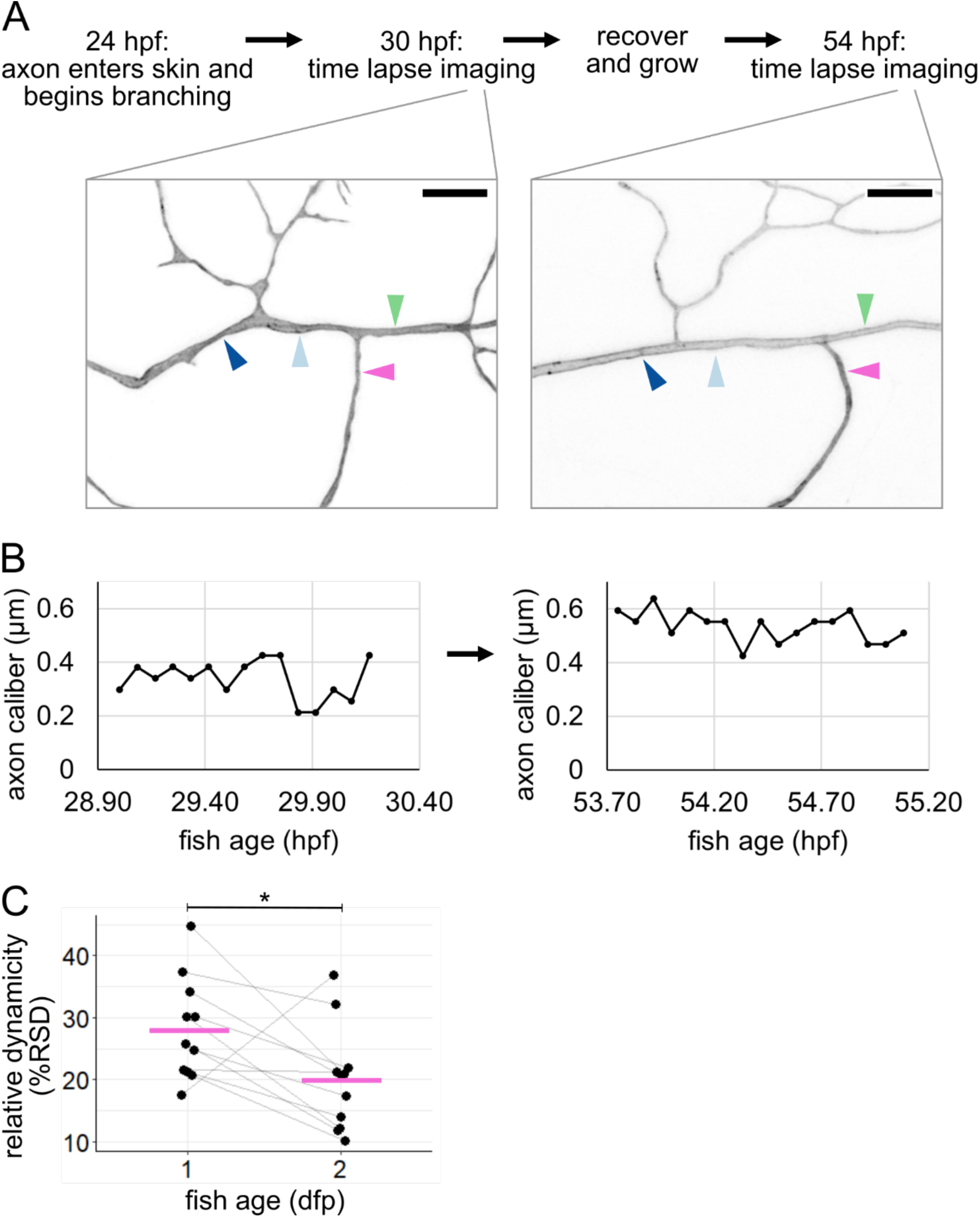
Tracking caliber dynamics over development **(A)** Experimental workflow with example images (same neuron as Figure 2). Arrowheads: locations for line scan measurements. Each location is 3 µm from a branch point and separated from other locations by at least one branch point. Scale bars: 5 µm. **(B)** Representative example plots of caliber dynamics for the same segment at 1 dpf and 2 dpf. **(C)** Plot of relative dynamicity (%RSD) at 1 dpf and 2 dpf (same neurons as Figure 5) n = 11 neurons, N = 5 fish. Grey lines connect data points from the same neuron. Magenta lines: mean values. *p < 0.05.

## Results

### Rohon-Beard (RB) neurons display a variety of calibers

RB neurons are touch-sensing neurons that differentiate and become functional within two days of development in embryonic zebrafish and other anamniotes (Katz et al., 2021). The peripheral axons of RB neurons can be identified by transmission electron microscopy (TEM) as membrane-bound ovals between the outermost (periderm) skin cells and lower (basal) skin cells in cross section (O’Brien et al., 2012). Using existing TEM data sets (Faas et al., 2012; O’Brien et al., 2012), we identified peripheral axons of RB neurons and trigeminal neurons, closely-related sensory neurons that perform the same touch-sensing function in the head. By measuring the short axis of these axon cross-sections, we found that calibers ranged from approximately 0.084 to 1.459 µm. This wide range prompted us to ask if axon caliber varies within each cell or only between cells, perhaps by sensory subtype (Gau et al., 2013; Palanca et al., 2013; Tuttle et al., 2024).

To determine if axon caliber can vary within an axon arbor, we sought to measure caliber at consistent locations across many intact cells. To enable absolute measurements of axon caliber, we developed an imaging protocol that paired sparse labeling with super-resolution microscopy. Using transient transgenesis, we generated embryos that expressed a membrane-localized fluorescent protein (EGFP-CAAX) in a single RB neuron in the tail (Fig 1B). We performed time lapse confocal microscopy starting shortly after RB neuron differentiation (20 hours post-fertilization, hpf) and confirmed that branches of the peripheral axon arbor form by growth cone bifurcation as well as collateral sprouting (Haynes et al., 2022; Sagasti et al., 2005), resulting in a highly branched axon arbor within hours of skin entry. To identify a consistent area of measurement, we observed the most proximal branch point in the peripheral arbor at 27 - 30 hpf and found that it typically forms by collateral sprouting (10 out of 10 cells, Fig 1C-C’, Video 1). In these images, some branches of each arbor appeared to be thick while others appeared to be thin (Fig 1D). This observation suggested that axon caliber can vary within each cell and supports the use of this location and technique for studying axon caliber variation.

### Caliber varies along the length of an axon segment

Previous studies have shown that axons are not cylindrical but vary in caliber along their lengths (Bar-Ziv et al., 1999; Greenberg et al., 1990; Griswold et al., 2024; Pullarkat et al., 2006). This phenomenon is sometimes referred to as pearling or varicose morphology. To determine if pearling is observed in live animals, we labeled individual RB neurons and measured axon caliber at 1-µm increments along the length of a segment, shortly after arbor formation began (29 hpf, Fig 2A-B). Variations in axon caliber were visible along the length of all resolvable axon segments, as has been described in mammalian neurons (Greenberg et al., 1990; Griswold et al., 2024). In some instances, axons had distinct pearls (Fig 2A’) and in others, more subtle thick and thin stretches without distinct inflection points (Fig 2C). These observations of lengthwise caliber variation confirm that fish and mammalian neurons share similarities in local axon morphology in vivo.

### Sister axon branch calibers are largely uncorrelated with each other

Next, we asked how axon caliber varies between different axon segments within a single cell. We labeled individual RB neurons and measured axon caliber near the branch point most proximal to the cell body in 27 - 31 hpf embryos (Fig 3A). The axon segment coming from the spinal cord was designated the primary axon (P), and the axon segments distal to the branch point were designated secondary axons (sister branches designated S1 and S2, for the thicker and thinner segments, respectively). Each segment was measured at three locations (3, 4, and 5 µm from the branchpoint), which were averaged to estimate the caliber of that axon segment. By measuring these locations, we observed that axon segments of different calibers exist within the same RB neuron and that secondary branches were typically thinner than the primary branch from which they arose (Fig 3B, P = 0.43 µm, S1 = 0.32 µm, S2 = 0.20 µm). These data confirmed that the average caliber of axons is not uniform across the cell.

Because sister segments, S1 and S2, often have different calibers, we sought to assess the relationship between them. We considered two hypotheses: 1) despite variations, each neuron has characteristically thick or thin calibers, and thus S1 and S2 have positively correlated calibers, or 2) S1 and S2 compete for cellular material, and thus their calibers are negatively correlated. To distinguish between these two hypotheses, we compared the S1 and S2 caliber for each neuron. Direct comparison showed a weak relationship between S1 and S2 caliber (Fig 3C, R^2^ = 0.13). This weak correlation suggests that neither hypothesis is strongly supported.

In keeping with this interpretation, symmetry across the branch point was weakly correlated with normalized S1 caliber (Fig 3D, R^2^ = 0.04), which we would expect to be strongly correlated if branches compete for cellular material to increase caliber. On the other hand, symmetry is more strongly correlated with normalized S2 caliber (Fig 3E, R^2^ = 0.54), suggesting that symmetry is achieved by the thinner branch being thicker, unrelated to the caliber of the S1 branch.

### RB axons taper

For an RB neuron to function properly, the cell needs electrical signals to travel efficiently. At the time points measured in Fig 3, the arbor was just a few hours old, functional, and continuing to expand. Already, optimal RB neuron function of transmitting sensory signals from the periphery requires efficient propagation of electrical signals from distal to proximal axon segments (toward the soma). Although the peripheral arbor of RB neurons transmits these afferent signals, as dendrites typically do, we refer to the peripheral arbor as an axon due to its uniformly plus-end-out microtubule organization (Shorey et al., 2021). Tapering is often considered a characteristic unique to dendrites (Craig and Banker, 1994; Jan and Jan, 2010), though more recent analyses suggest that branched axons may taper as well (Desai-Chowdhry et al., 2022).

To facilitate comparison of RB axon branching to other systems, we assessed tapering. First, we calculated the cross-sectional area of each primary and secondary branch (area = 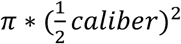 We compared the primary cross-sectional area to the sum of the secondary cross-sectional areas (Fig 4A). On average, the ratio of the combined secondary cross-sectional areas to the primary cross-sectional area was less than 1 (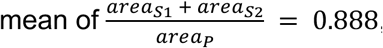, Fig 4B), meaning that the RB peripheral arbor tapers in area.

Tapering can influence the health and function of a neuron in many ways, some of which scale with area, and some of which scale with radius (Craig and Banker, 1994; Desai-Chowdhry et al., 2022). Hence, we also compared the radius scaling ratio between secondary branches and their corresponding primary branches (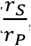, Fig 4C). The average ratio for the thicker branch was much larger than for the thinner sister branch (mean 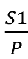 = 0.77 ± 0.033 𝑆𝐸𝑀, and mean 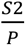 49 ± 0.030 𝑆𝐸𝑀, Fig 4D). In evaluating a branched system, some scaling rules hold for the population average, even when branches are asymmetric (Brummer et al., 2017). Hence, to evaluate the function of the arbor more generally, we combined the measurements for the two branches and found that the average ratio (mean 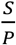= 0.629 ± 0.0282 𝑆𝐸𝑀) was similar to what is predicted for a system that optimizes for power dissipation rather than time delay (Desai-Chowdhry et al., 2022).

Here, we refer to the RB peripheral arbor as an axon due to its plus-end-out microtubule organization (Shorey et al., 2021). Therefore, our observations demonstrate that it is possible for neuronal processes with an axon-like cytoskeletal organization to taper.

### Axon caliber is highly dynamic

Having documented variability in axon caliber across locations within each RB neuron, we wondered how calibers change with time. To investigate caliber dynamics, we performed time-lapse imaging in the proximal region of the tail-innervating axon arbors at 28 - 31 hpf (Video 2). Axon caliber changed in several qualitatively different ways over time. We categorized these dynamic behaviors into four categories (Video 3): (1) a traveling pearl, instances of bubble-like regions moving along an axon (Fig 5A), (2) focal inflation and deflation, in which a pearl appears and disappears over the course of multiple time points (Fig 5B), (3) segment widening and narrowing, in which an entire segment thickens and thins between time points (Fig 5C), and (4) a constriction point that appears and disappears, sometimes repeatedly (Fig 5D). This variety of dynamic behaviors suggests that several distinct processes likely contribute to axon caliber dynamics.

Axon caliber is known to be altered by changes in target cells (Nabel et al., 2024; Sinclair et al., 2017; Voyvodic, 1989a) or myelination (Ciocanel et al., 2020; Seidl, 2014; Walker et al., 2019), but these developmental processes occur on different timescales from the dynamics we observed with intervals of 5 minutes (Fig 5A-D). To assess caliber dynamics quantitatively, we selected several locations within the proximal region of each neuron, each 3 µm away from a branch point, and measured axon caliber at that location over the course of imaging (Fig 5E). Axon caliber was highly dynamic (Fig 5F), with standard deviations ranging from 0.045 µm to 0.095 µm, and could vary with a range of up to four-fold at the same location in less than two hours. Hence, axon caliber and local morphology may be more dynamic than has been widely appreciated.

Next, we asked if thin or thick axon segments are more dynamic. First, we assessed absolute dynamicity by comparing average caliber to the standard deviation (SD) or measurements observed during the time lapse (Fig 5G). We found that SD and axon caliber correlated positively, suggesting that thicker axons experienced larger or more fluctuations in caliber. A possible caveat for our measurements is that the lower dynamicity could be due to thin axons spending more time below the limit of resolution. To determine if this was the case, we also assessed relative dynamicity. We calculated relative standard deviation (RSD) for each location by dividing the standard deviation by the average caliber across the movie. Comparing dynamicity across segments revealed that caliber and RSD were negatively correlated, meaning that thick axons were less dynamic than thin axons (Fig 5H). If differences in absolute dynamicity were due to resolution limitations, we would expect to see that relative dynamicity of the thinnest axons is also low. Instead, thin axons had high relative dynamicity, indicating that, if anything, we may have underestimated the dynamicity of thin axons. Together, these results suggest that thick axons have more or larger absolute fluctuations in caliber, but the fluctuations experienced by thin axons are more dramatic when compared to their average caliber.

### Caliber dynamicity changes over development

Because we observed caliber to be highly dynamic early in the life and function of RB neurons, we sought to determine if caliber continues to be dynamic later in development. To do this, we recovered embryos after imaging at 28 - 31 hpf and allowed them to continue developing for an additional 24 hours. These fish were re-mounted at 52 - 55.5 hpf and imaged by time-lapse microscopy again. Although all cells had changed shape and axon arbors had grown, often the locations imaged on the first day could be identified, imaged, and measured again on the second day (Fig 6A-B). We found that all resolvable axons continued to be dynamic. However, dynamicity measured as %RSD decreased from the first to the second day in most axon segments (10 out of 11 axon segments decreased. Mean %RSD = 28% at 1 dpf and 20% at 2 dpf, p = 0.036, Fig 6C). Because some axon segments got thicker (n = 6 out of 11) and some got thinner (n = 5 out of 11), this decline in relative dynamicity is not simply a product of axons becoming universally thicker. Instead, this dynamicity likely reflects developmental changes that occur in the cell and tissue.

### Cellular microenvironment can impact axon caliber

Based on the qualitative variety of caliber dynamics we observed on the minutes timescale (Fig 5A-D), we hypothesized that there are multiple contributors to caliber dynamics. Cell-intrinsic effectors that may cause axon caliber to be dynamic on short timescales, such as cargo transport and contraction of the membrane periodic skeleton, have been described by others (Costa et al., 2020; Wang et al., 2020). However, another contributor is the cellular microenvironment. Mechanical force at the soma has been shown to influence axon and dendrite morphology (Pan et al., 2024). We hypothesized that, in RB peripheral axons, caliber changes could be influenced locally, including by contact with surrounding epithelial cells.

Because RB axons are embedded in actively developing skin, they are subject to local deformation caused by morphological changes in surrounding epithelial cells. Specifically, RB peripheral axons grow between two epithelial cell layers, the periderm and basal cells, each of which undergoes rapid expansion at these developmental stages (Fig 1A) (O’Brien et al., 2012).

To determine if changes to the shape or adhesion of adjacent epithelial cells can impact the caliber of axons, we performed simultaneous time lapse imaging of axons and the underlying basal cells starting at 28 - 31 hpf (Video 4). We imaged basal cells using a marker of lipid microdomains under the control of the ΔNp63 promoter (Rasmussen et al., 2015; Rosa et al., 2023). One of the most dramatic and frequent perturbations to the RB axon environment occurs when basal cells round up for mitosis, during which the lipid reporter clearly highlights rounded cells against the field of surrounding, flat basal cells (Fig 7A). We measured caliber at 1-µm intervals along the length of the axon, from a flat neighbor cell, onto the rounded cell, and onto the next flat neighbor cell (Fig 7B). We found that segments of axon on the rounded cell were significantly thicker than segments of the same axon on the non-dividing neighbor cells (caliber = 0.21 µm on rounded cell, 0.16 and 0.14 µm on non-dividing neighbors, p-value = 0.025 and 0.00035, respectively, n = 114 measurement locations, N = 4 axons/fish, Fig 7C). This difference contrasts with the two non-dividing neighbor groups, which did not have significantly different calibers, as expected (p-value = 0.059). Although this observation does not distinguish between mechanical or signaling interactions as the driving factor for caliber changes, these data align with a model in which dynamic properties of the dividing cell may impact local axon caliber.

**Figure 7:**
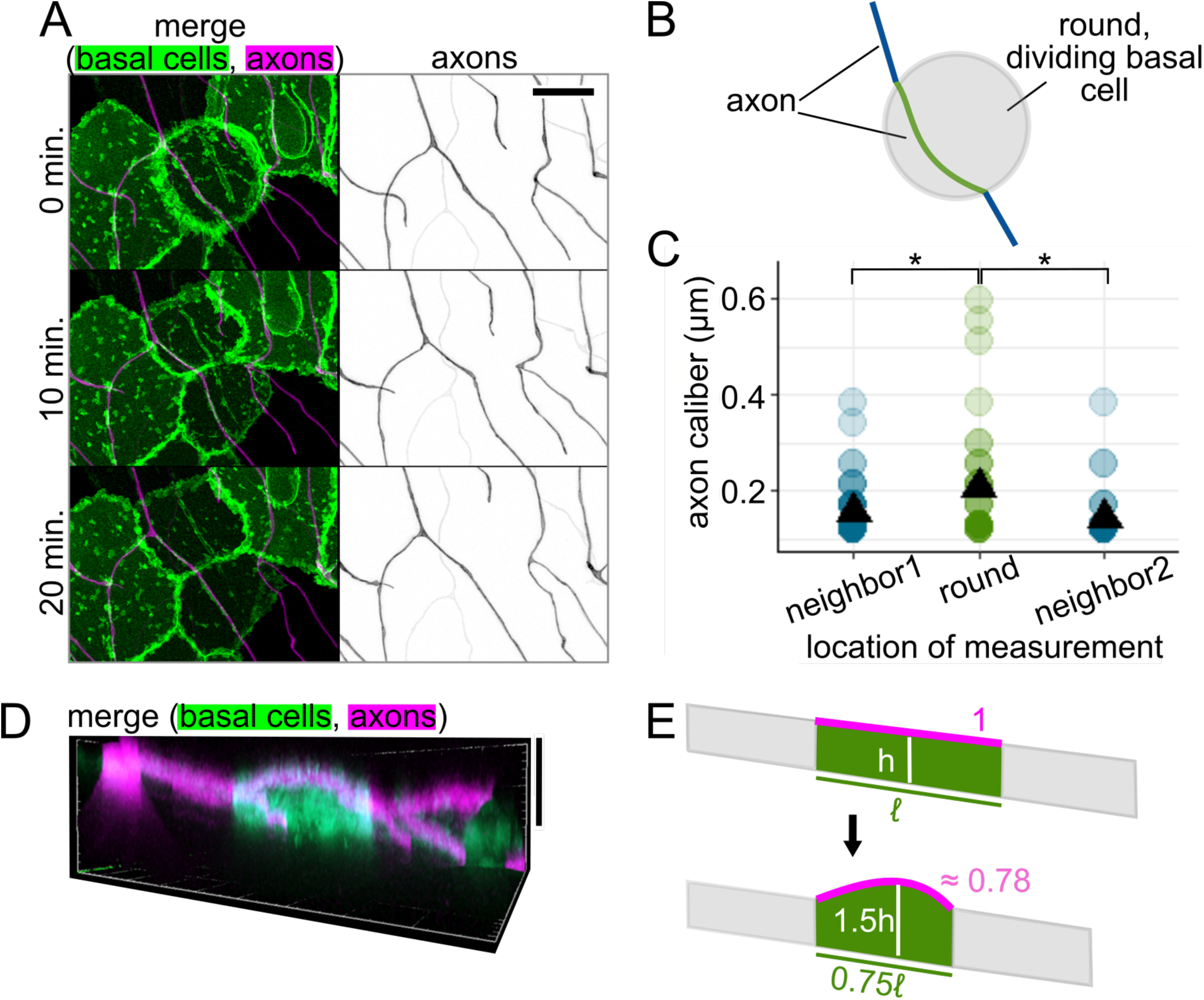
Axon caliber on dividing basal epithelial cells **(A)** Representative images of time lapse showing an RB axon adjacent to a dividing basal cell. Basal cells express EGFP-PLCδ-PH. RB neurons express mRuby-CAAX. Scale bar: 10 µm. **(B)** Diagram of an axon on a dividing basal cell that is rounded while undergoing mitosis. Green line: axon segment in contact with dividing basal cell. Blue lines: axon segments on neighboring, non-dividing basal cells. **(C)** Plot of axon caliber measurements grouped by location, n = 114 measurement locations, N = 4 axons/fish. Green dots: caliber of axon on a dividing basal cell. Blue dots: caliber of axon on neighboring cells to one side (neighbor1) or the other (neighbor2) of the dividing basal cell. Black triangles: mean value. *p-val < 0.05. **(D)** Orthogonal projection of a dividing basal cell, which expresses EGFP-PLCδ-PH and is highlighted against non-dividing basal cells due to a variegated expression pattern. An RB axon is in contact with its apical surface, which is rounded upward. Vertical scale bar: 10 µm. **(E)** Diagram comparing axon path length on a rounded, dividing basal cell to its flattened state as an approximation of how basal cell shape changes may impact tension on the axon. Magenta: approximated, theoretical axon length, estimated as the longest distance across the apical surface in an orthogonal projection. Approximate arc length was calculated based on height and length estimates in orthogonal projections. ℓ: original length of basal cell, h: original height of basal cell.

### Forces from surrounding cells may impact axon morphology

One reason why axons might be thicker on rounded cells could be due to changes in tension (Griswold et al., 2024; Pan et al., 2024). As a basal cell enters mitosis, it pulls its borders inward, becoming shorter in planar distance, and rounds up, creating a greater vertical distance (Fig 7D). We used images with the clearest orthogonal projections to obtain estimates of how the dimensions of a dividing basal cell differs from its non-dividing neighbors. Using these estimates, we calculated theoretical changes in axon length as the cell rounds up (Fig 7E). To simplify these calculations, we assumed that the axon is fixed at the cell borders and runs in a straight path across the basal cell surface. We found that the rounding of basal cells resulted in a shorter axon path during division than when cells were flat (path length when round ≈ 0.78 * path length when flat). Assuming that axons are adhered to the non-dividing neighboring cells, these estimates are consistent with the axon being under lower tension on the rounded cell than on a non-dividing neighbor cell.

Based on our observations of rounded cells and our estimates of length changes, we expected that axon caliber would decline as the daughter basal cells flatten. To test this prediction, we performed time-lapse imaging with 5-minute intervals on the rounded cells, including those in Fig 7C, to see if axon caliber changes as the rounded cell divides and flattens into two daughter cells. For each time point, we measured the distance between the two points where the axon crossed the border of the dividing cell or its daughter cells (Fig 8A). The two time points between which the change in length was greatest were selected, and the caliber of the axon on the rounded cell or its daughters was measured at 1-µm increments for those two time points (Fig 8B-C). We paired measurements by location to assess if there was a significant drop in axon caliber between these two time points. Although axon caliber declined from the round to flat time point, on average, this subtle decrease did not reach statistical significance (caliber = 0.28 µm when the basal cell was round and 0.25 µm when the basal cell was flat, p-value = 0.17, n = 54 measurements, N = 7 axons/fish, Fig 8D). This contrasts with the portions of the axon that were on non-dividing neighbor cells, where the difference could have easily occurred by random chance (p-value = 0.94, n = 37 measurements). This result suggests that the decrease in average axon caliber as the underlying basal cell flattened is small and likely not meaningful.

**Figure 8:**
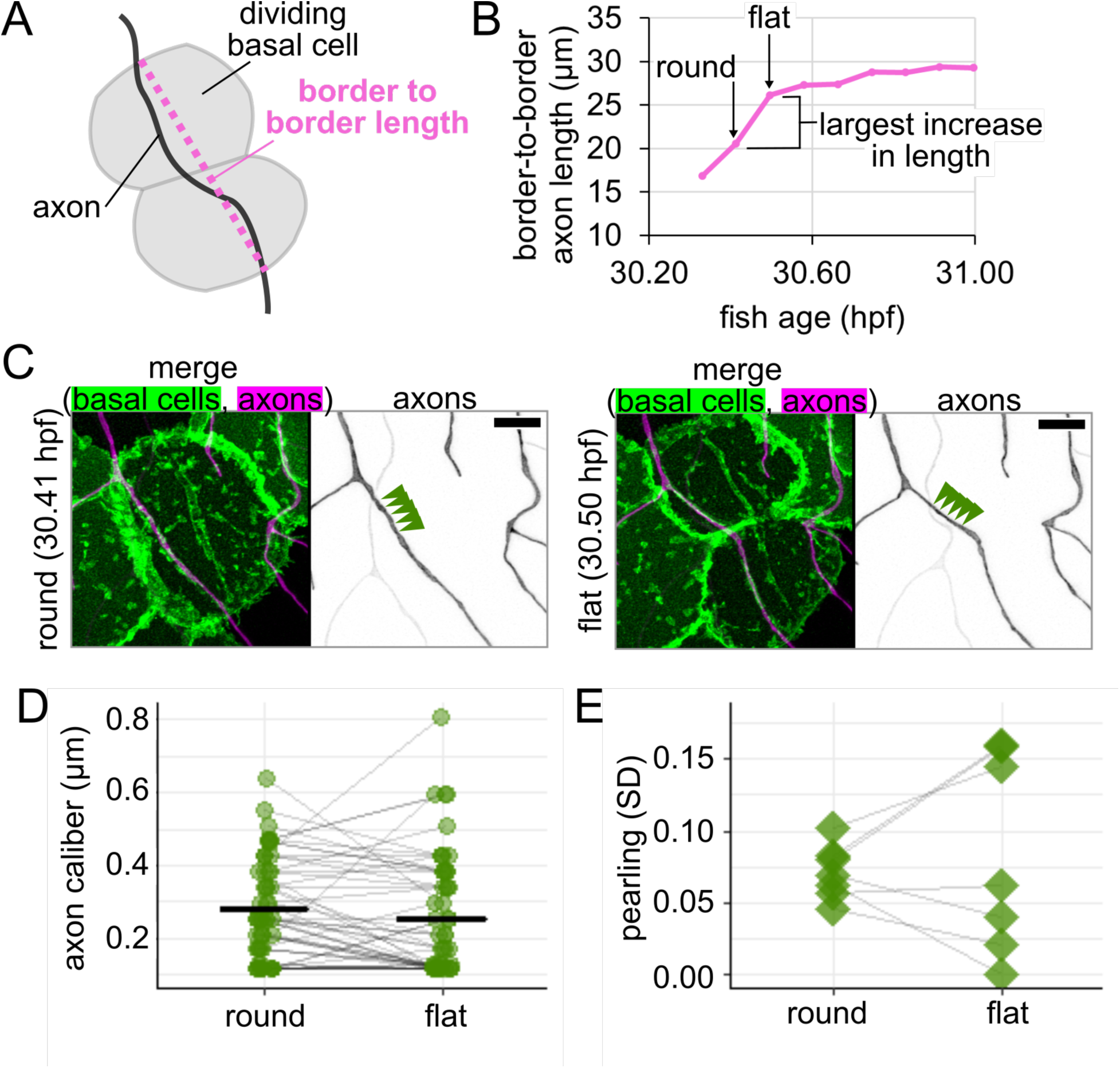
Changes to axon morphology as new basal epithelial cells flatten **(A)** Diagram of how border-to-border length was measured for an axon on a dividing basal cell. **(B)** Example plot of changes in border-to-border axon length during time lapse imaging for axon shown in Fig 7A. Round and flat time points are defined as sequential points (5-min. interval) during which border-to-border length increases the most. **(C)** Images from time points highlighted in panel B. Arrowheads: five measurement locations. Scale bars: 5 µm. **(D)** Plot of axon caliber for all locations that remained on the dividing basal cell or its daughter cells, n = 54 measurement locations, N = 7 axons/fish. Grey lines connect data points from the same location. Black lines: mean values. **(E)** Plot of pearling, defined as SD across all locations, to assess morphology change for axons shown in panel D, N = 7 axons/fish. Grey lines connect data points from the same axon.

Despite the lack of change in average caliber, visual assessment of the videos appeared to show a change in morphology during basal cell flattening (Fig 7A, 8C). Tension and mechanical force may be an important parameter in pearling (Bar-Ziv et al., 1999; Griswold et al., 2024; Pan et al., 2024; Pullarkat et al., 2006), so we wondered if thick regions (i.e. pearls) and thin regions (i.e. connectors) undergo different changes during basal cell division.

To assess morphological changes to axons during basal cell division, we calculated pearling as the standard deviation of measurements taken along the length of each axon for the rounded time point and for the flat time point. By pairing values from each movie, we found that one subset of axons became more pearled (increased SD) and the rest became less pearled (decreased SD), resulting in two groups of cellular responses to flattening (Fig 8E). This heterogeneity might suggest that either 1) different responses arise from heterogeneity among the RB neuron population, or 2) our 5-minute intervals do not provide sufficient time resolution to capture cellular responses to tension, which may occur in less than 10 minutes (Sinha et al., 2011; Sitarska and Diz-Muñoz, 2020). Regardless of the reason, these results demonstrate that axon morphology changes with basal cell flattening, though additional complexities remain to be explored.

## Discussion

### Tapered architecture in branched sensory axons

We have found that the calibers of axon segments in the same RB neuron can be largely independent of one another, highly dynamic, and likely shaped by multiple determinants, including contact with epithelial cells. Variation in axon caliber had previously been observed along short stretches of fixed axons in rodent neurons (Greenberg et al., 1990; Griswold et al., 2024), in response to developmental changes to the neuron’s environment and signaling (Bin et al., 2024; Ford et al., 2015; Kumar et al., 2005; Metzner et al., 2022; Nabel et al., 2024; Sánchez et al., 1996; Seidl, 2014; Seidl et al., 2010; Seidl and Rubel, 2016; Sinclair et al., 2017), and with forces on millimeters-length scales (Pan et al., 2024). Our results expand upon these findings by characterizing caliber variations in vivo across a branched axon, with fine timescale, and with local deformations caused by epithelial cell division.

The cutaneous axon endings of touch-sensing neurons, including trigeminal, dorsal root ganglion (DRG), and RB neurons, share characteristics with both dendrites and axons. Like dendrites, they detect stimuli from the periphery and relay signals towards the cell body, but, like axons, their microtubules are oriented plus-end-out (Shorey et al., 2021). We found that the branched axon arbors of RB neurons taper across the most proximal branch point, becoming narrower in caliber and cross-sectional area further from the soma (Fig 4). This feature is often regarded as a characteristic of dendrites, not of axons (Craig and Banker, 1994; Jan and Jan, 2010). Mechanisms of signal relay from periphery to soma in RB neurons have not been fully elucidated. If RB neurons fire action potentials, then their tapering (Fig 5) would favor afferent action potential conduction, as signals travel from the periphery toward the soma. In measuring the radius scaling ratio, we found it to be consistent with neurons optimized for power dissipation, which aligns well with a more graded, dendrite-like signal transduction (Desai-Chowdhry et al., 2022). Together, our findings are consistent with a model in which tapering may be most aligned with the direction of functional signal propagation rather than microtubule polarity. Whether our findings hold true for other axons, especially afferent axons, remains to be explored.

In comparing sister axon branches, we hypothesized that sister branches are related to each other, resulting from either cell-wide effects or a competition for cellular materials. Instead, we found that their calibers are largely uncorrelated with each other, despite being in close proximity and sharing a cytoplasm. Interestingly, we observed varying degrees of symmetry, ranging from pairs of sister branches with the same caliber to highly asymmetrical pairs, where one secondary branch is much thicker than the other. One reasonable hypothesis to explain this observation could be that branches of different symmetry form by different mechanisms. For example, highly asymmetric branches might have formed by collateral sprouting and symmetric branches by growth bifurcation. However, we found a full spectrum of symmetry values, not a bimodal distribution, arguing against this hypothesis (Fig 3D-E). Moreover, though bifurcations are common in RB arbor outgrowth (Haynes et al., 2022) the first branch point often, if not always, forms as a collateral (Fig 1C). Thus, collateral sprouting can give rise to segments of a variety of calibers within a few hours of formation, highlighting the malleability of axon caliber.

### Mechanisms of caliber variation over time and space

We found that caliber dynamicity changed over development (Fig 6). Many developmental changes could account for a shift in dynamicity, including changes within the cell itself, such as cargo transport (Costa et al., 2020; Wang et al., 2020), or changes to the extracellular environment, such as axon ensheathment (Jiang et al., 2019; O’Brien et al., 2012; Rosa et al., 2023). Time lapse imaging of axon caliber also revealed multiple qualitatively different types of dynamics (Fig 5A-D), which we refer to as traveling pearls, focal inflation and deflation, segment widening and narrowing, and constriction points. This variety suggests that there are likely multiple contributors to variations in axon caliber, including both cell-intrinsic and cell-extrinsic mechanisms.

Cell-intrinsic determinants of axon caliber, such as the cytoskeleton, have been studied to varying degrees. One of the best-known caliber determinants is neurofilaments, a type of intermediate filament specific to the nervous system. Increased neurofilament abundance increases axon caliber in mammalian and bird neurons (Elder et al., 1998; Eyer and Peterson, 1994; Kriz et al., 2000; Ohara et al., 1993; Sainio et al., 2021; Sakaguchi et al., 1993; Zhu et al., 1997). Neurofilaments are also highly dynamic. Differences in transport, differential modifications, and dynamics (e.g. folding, severing, annealing) of neurofilaments are all possible ways in which these filaments might contribute to both local regulation and axon caliber dynamics within a cell (Boyer et al., 2022; Çolakoğlu and Brown, 2009; Uchida et al., 2023).

Another cytoskeletal candidate for cell-intrinsic effectors of caliber variation is the more recently discovered membrane periodic skeleton (MPS), regularly-spaced actin rings connected by spectrin tetramers that run the length of axons and dendrites (He et al., 2016; Xu et al., 2013). These rings expand when large cargo such as mitochondria and lysosomes pass through, and altering myosin contractility changes caliber and retrograde trafficking (Costa et al., 2020; Leite et al., 2016; Wang et al., 2020). Observations of cargo transport through the MPS (Costa et al., 2020; Wang et al., 2020), as well as fixed observations that large cargo is often found within pearls (Greenberg et al., 1990), suggest that intracellular cargo transport along microtubules may also contribute to axon caliber dynamics. Work by Griswold et al. demonstrated that myosin contractility can impact pearling (Griswold et al., 2024). Furthermore, the MPS and the propensity of neurons to bead have been found to be protective against low-level mechanical forces (Krieg et al., 2017; Pan et al., 2024), highlighting MPS contractility as another potential source of caliber variations.

Other intracellular factors, such as importin beta (Bin 2024), PI3K/AKT signaling (Kumar 2005) and membrane composition (Griswold 2024) also regulate axon caliber. Although some of these factors likely influence caliber by regulating the cytoskeleton (Bin 2024), membrane composition is a non-cytoskeletal, cell-intrinsic candidate for causing caliber variation. Griswold et al. found that membrane fluidity can change which size and shape of pearls are most energetically favorable, thereby changing axon morphology (Griswold et al., 2024). Thus, given the dynamic nature of caliber in vivo, changes to membrane composition could also change caliber dynamics.

Damage can also influence axon morphology. Markedly pearled or varicose morphologies, sometimes referred to as beading or swelling, often form in damaged axons (Datar et al., 2019; Kerschensteiner et al., 2005; Shao et al., 2020; Williams et al., 2014). Indeed, damaged trigeminal and RB axons, themselves, become dramatically more beaded just before degeneration (Martin et al., 2010). Mechanical forces below the level necessary to break axons have also been found to impact axon and dendrite pearling (Pan et al., 2024). Our observations of variation in caliber in this study were made in anesthetized, healthy embryos and are unlikely to result from axon damage, as neurons remained intact, even after time lapse microscopy. Similarity between these phenomena in healthy and damaged axons may reflect related processes if the healthy malleability of axon caliber is pushed to extremes upon insult.

Although much attention has focused on potential intrinsic determinants for axon caliber variation, we considered the possibility that extrinsic influences from surrounding cells might affect axon caliber locally. Cell-extrinsic factors that influence axon caliber include target size (Voyvodic, 1989a) and distance (Seidl et al., 2010), neuronal function (Nabel et al., 2024; Sinclair et al., 2017), and mechanical forces at millimeters-length scale (Pan et al., 2024). Each of these factors likely impacts the entire cell and is, therefore, unlikely to regulate caliber variation locally.

Our study focused on smaller deformations that occur on smaller length scales and timescales. Although myelination is thought to influence local axon caliber (Rydmark, 1981; Seidl, 2014; Swärd et al., 1995), questions remain about the direction and specific nature of this intercellular relationship (Bin et al., 2025; Friede, 1972; Sánchez et al., 1996). RB axon arbors are embedded in a developing epidermis, which undergoes growth and morphogenetic changes. RB axons span tens or even hundreds of micrometers in length and branch to cover large areas of the skin. Hence, each segment of an axon contacts different basal cells, which may each impact the local axon caliber independently.

Basal cells can undergo many morphological changes to alter the axon’s environment. One important change is ensheathment (O’Brien et al., 2012) (Fig 1A), which increases between 1 and 2 dpf (Jiang et al., 2019; Rosa et al., 2023), possibly contributing to differences in dynamicity between these two developmental time points (Fig 6C). The idea that ensheathment may alter axon caliber is further consistent with the observation that myelinated portions of an axon are thicker than the nodes of Ranvier (Rydmark, 1981; Swärd et al., 1995).

Another morphological change in basal cells is rounding for division, which we found changes axon caliber and morphology (Fig 7,8). We hypothesize that changes in tension caused by the movement of cells adjacent to axons are candidates for cell-extrinsic regulators of axon caliber. Division events likely vary in frequency throughout embryonic development, which may further contribute to developmental differences in caliber dynamicity (Fig 6C).

Additionally, axons may experience end-to-end stretch as embryos lengthen. Finally, although our studies were in embryonic zebrafish, when the environment surrounding these neurons is a rapidly expanding bilayered epidermis, dynamics are unlikely to cease in adulthood, since the stratified epidermis in both fish and mammals is constantly renewing as basal cells differentiate into higher strata. How caliber variations and dynamics differ in this stratified system remains to be explored. In conjunction with findings that myelination influences axon caliber (Ciocanel et al., 2020; Rydmark, 1981; Seidl, 2014; Swärd et al., 1995; Walker et al., 2019), our results support the model that lengthwise variation, local variations in average caliber, and caliber dynamics may be influenced, in part, by the behavior of surrounding cells.

## Video Legends

**Video 1: Time-lapse microscopy of axon branch formation in RB neurons.**

Confocal time lapse microscopy of RB neuron development in the embryonic zebrafish tail. The main branch of the peripheral axon arbor grows out of the developing spinal cord and into the skin. Shortly thereafter, many branches form by growth cone bifurcation and collateral sprouting. Pseudocoloring in the final frames highlights proximal arbor, and arrows indicate branch points that formed by collateral sprouting events at the times indicated. Scale bars: 5 µm.

Video 2: Time-lapse microscopy of peripheral RB axon.

Super-resolution time-lapse microscopy of the peripheral axon arbor showing the most proximal branch points. Frequent and rapid changes in axon caliber occur within the approximately 1 hour captured prior to drift. Time intervals: 5 minutes. Scale bar: 5 µm.

Video 3: Types of caliber fluctuations in RB axons.

Super-resolution time-lapse microscopy of peripheral axon arbors showing examples of qualitatively different fluctuations in axon caliber. Time intervals: 5 minutes. Scale bars: 2 µm.

**Video 4: Time-lapse microscopy of RB axon in contact with mitotic basal keratinocyte.** Super-resolution time-lapse microscopy of an RB axon segment in contact with a skin cell that is undergoing mitosis. Morphological changes in both cells were visualized via membrane fluorescent membrane tags. Magenta: RB neuron labeled with mRuby-caax. Green: Basal keratinocyte labeled with egfp-plcδ-ph. Time interval: 5 minutes. Scale bar: 5 µm.

## Acknowledgements

The studies were supported by National Institutes of Health grant R01AR064582. The content is solely the responsibility of the authors and does not necessarily represent the official views of the National Institutes of Health. We thank Van Savage for valuable discussions and suggestions regarding branch analysis, Melissa Emami for providing the Isl1[ss]:lexA, lexAop:mRuby-caax fish, D’Juan Farmer and the UCLA Broad Stem Cell Research Center Microscopy Core for microscope access, and the UCLA Zebrafish Core Facility for animal care. We thank the Sagasti, Farmer, and Koehler labs for helpful discussion and suggestions.

